# Ascending propriospinal modulation of thoracic sympathetic preganglionic neurons during lumbar locomotor activity

**DOI:** 10.1101/2025.10.14.681171

**Authors:** Lucia E Dominguez-Rodriguez, Chioma V Nwachukwu, Narjes Shahsavani, Juanita Garcia, Jeremy W Chopek, Kristine C Cowley

## Abstract

Although the autonomic sympathetic system is activated in parallel with locomotion, the underlying neural mechanisms mediating this coordination are not completely understood. Descending exercise or ‘central command’ signals from hypothalamic and brainstem regions are thought to activate thoracic spinal sympathetic neurons in parallel with descending locomotor commands. In turn, subsets of thoracic sympathetic preganglionic neurons (SPNs) then increase activation of a constellation of tissues and organs that provide homeostatic and metabolic support during movement and exercise. It is known that neurons within the spinal cord (propriospinal networks) can generate well-coordinated and sustained locomotor activity but whether these propriospinal networks contribute to coordination between locomotor and autonomic systems is unknown. To investigate this, we applied neurochemicals to elicit whole-cord or lumbar-evoked locomotor activity in an *in vitro* spinal cord preparation, simultaneously recording lumbar ventral root (VR) activity and changes in calcium fluorescence of pre-labelled SPNs in thoracic segments. Using whole-bath drug application to elicit hindlimb locomotor activity, recorded SPN responses were increased in rostral (T4 – T7) compared to caudal (T8 – T11) segments. When locomotor-inducing neurochemicals were applied only to the lumbar region using a split-bath configuration, SPN population responses were increased in rostral (T4-7) but not caudal (T8-9) segments during both tonic and rhythmic VR activity. In both approaches, the greatest numbers of SPNs with increased fluorescence during rhythmic activity were in T6/7, whereas the greatest numbers with unchanged or decreased fluorescence were in caudal segments (T8-T11). Together these findings reveal a strong ascending lumbar to thoracic integrating communication pathway and may represent a key feature of spinal neural network function normally. Such communication pathways should be further investigated for targeted autonomic function(s) activation and therapeutic benefit after spinal cord injury.

## INTRODUCTION

Locomotion and exercise are fundamental behaviors involving sustained rhythmic contractions of muscles, at varying intensities, depending upon the mode of movement and groups of muscles used. It is well established that the generation of a locomotor pattern for overground movement is an intrinsic property of the spinal cord, with important contributions from afferent and descending supraspinal inputs [reviewed in (Stuart and Hultborn 2008)]. For generating basic patterns of rhythmic flexion and extension, the motor cortex and basal ganglia generate and select motor commands, which are integrated in the mesencephalic locomotor region (MLR). Projections from the MLR convey information to regions in the medullary reticular formation (medRF) which acts as a final integration and relay centre for descending locomotor command signals. These descending command signals activate spinal neural circuitry which converts tonic descending drive command signals into rhythmic and well-coordinated locomotor activity, in spinal networks known as locomotor or central pattern generator networks (i.e., LPGs, CPGs) (Shik, Severin et al. 1969, Jordan, Pratt et al. 1979, Grillner 1981, Whelan 1996, Jordan, Liu et al. 2008). Electrophysiological and pharmacological evidence supports a fast glutamatergic and slower serotonergic locomotor pathway from MLR to medRF and to spinal locomotor networks (Liu and Jordan 2005, Jordan, Liu et al. 2008).

Electrical stimulation of the MLR to induce locomotion also causes concomitant increases in blood pressure (Eldridge, Millhorn et al. 1985, Bedford, Loi et al. 1992), in part mediated by descending input to spinal sympathetic networks via vasomotor commands from the rostro-ventral lateral medulla (RVLM) ((Loewy and Spyer 1990, Schreihofer and Sved 2011). Indeed, the RVLM is a key integration site that regulates homeostatic and metabolic functions integral for maintaining whole body homeostasis (i.e., blood pressure, temperature, respiration and heart rate regulation) and for regulating availability of circulating glucose and fatty acids by regulating glucose counterregulatory responses and lipolysis from adipose tissue stores, respectively (Blessing, Yu et al. 1999, Morrison, Sved et al. 1999, Bartness, Shrestha et al. 2010, Bartness, Vaughan et al. 2010, Bartness, Liu et al. 2014, Verberne, Sabetghadam et al. 2014). However, the neural mechanisms underlying the integrated activation of locomotor and sympathetic autonomic systems are only recently starting to be understood. For example, a monosynaptic glutamatergic excitatory projection from neurons in the MLR to RVLM neurons was recently identified, and optical stimulation of these MLR neurons increased mean arterial pressure and elicited tonic discharge in L5 ventral roots in a decerebrate rat preparation (Koba, Kumada et al. 2022). Pulsed optical stimulation of MLR neurons in freely moving rats increased mean arterial pressure and speed of ongoing locomotion, suggesting locomotor and autonomic sympathetic systems were functionally integrated in intact preparations (Koba, Kumada et al. 2022). Similarly, focused electrical stimulation of the parapyramidal region in the medulla induces hindlimb locomotor activity in the *in vitro* neonatal rat (Liu and Jordan 2005) and chemogenetic and optogenetic activation of serotonergic neurons in the parapyramidal region increases both blood pressure and hindlimb locomotor activity in the adult decerebrate rat (Armstrong, Nazzal et al. 2017, Armstrong 2024). These blood pressure increases precede and outlast bouts of hindlimb locomotor activity.

Despite clinical observations that spinal electrical stimulation in lumbar locomotor regions can improve or normalize a subset of movement and exercise-related metabolic and homeostatic functions in motor complete spinal cord injury (reviewed in (Flett, Garcia et al. 2022)), less is known about whether and how locomotor and sympathetic systems are integrated at the spinal level.

Recently, we demonstrated that a class of genetically identified, locomotor-related neurons (V3 INs) in the lumbar region provide direct excitatory glutamatergic synaptic projections onto thoracic spinal preganglionic sympathetic neurons (SPNs), and that optical stimulation of lumbar V3 INs generates action potentials in SPNs (Chacon, Nwachukwu et al. 2023). Thus, at least one class of spinal locomotor-related neurons contribute to excitation of thoracic SPNs, demonstrating a neural substrate/mechanism of intraspinal ascending locomotor-sympathetic system integration.

Although SPN activity in response to application of glutamate, serotonin or dopamine agonists has been characterized in slice or spinal preparations (Spanswick and Logan 1990, Pickering, Spanswick et al. 1994, Gladwell and Coote 1999, Zimmerman, Sawchuk et al. 2012), SPN responses during rhythmic locomotor activity have not been described. Additionally, SPN responses to the locomotor activity-inducing combination of these neurotransmitters has not been described or characterized by segmental level. Thus, the present work examines the effect(s) of tonic and rhythmic locomotor activity on SPN responses at different thoracic rostro-caudal levels (T3/4 through T11).

The results provide physiological evidence that ascending propriospinal pathways contribute to lumbar locomotor-mediated activation of sympathetic neurons in particular thoracic segments, while decreasing activation of SPNs at other thoracic levels. Preliminary results were presented previously in abstract form (Nwachukwu, Chacon et al. 2022, Dominguez-Rodriguez, Nwachukwu et al. 2024).

## METHODS

### Animals

Experimental protocols used complied with the guidelines set by the Canadian Council on Animal Care and were approved by the University of Manitoba animal ethics committee. In our initial set of experiments, postnatal day 0 (P0) to P5 pups from our in-house C57Bl/6 colony were used. In all subsequent experiments, B6.129S-Chat^tm1(cre)Lowl^/MwarJ (Jax strain # 031661) were crossed with B6J.Cg-Gt(ROSA)26Sor^tm96(CAG-GCAMP6s)Hze^/MwarJ (Jax Strain # 028866) to generate B6.129S- Chattm1(cre0Lowl/Gt(ROSA)26Sortm96(CAG-GCamp6s)Hze/Mwar mice (referred to ChatCre/Gcamp6s). Pups (P0-P5) from the ChatCre/Gcamp6s endogenously express the genetically encoded calcium indicator (GCamp6s) in cholinergic SPNs and motoneurons (MNs) and were used for experimentation.

### *In vitro* electrophysiology

#### Whole spinal cord preparation with obliquely cut surface

Spinal cords were dissected from either C57BL/6 (n = 26) or ChAT-Cre/GCaMP6S (n = 30) mice, as previously described (Chopek, Nascimento et al. 2018, Chacon, Nwachukwu et al. 2023). Briefly, animals were anesthetized with isoflurane (Fresenius Kabi Canada Ltd, ON, Canada) decapitated at the medulla-spinal cord junction, eviscerated, and spinal cords dissected out in ice-cold (∼4°C) dissecting artificial cerebrospinal fluid (aCSF), composed of (mM): KCl (3.5), NaHCO_3_ (35), KH2PO_4_ (1.2), MgSO_4_ (1.3), CaCl_2_ (1.2), glucose (10), sucrose (212.5), and MgCl_2_ (2.2). The dissecting aCSF solution was equilibrated to pH 7.4 and continuously oxygenated (95% O2; 5% CO_2_). Spinal cords were then mounted on an agar block and fixed in place with acrylic glue for sectioning using a vibratome (Leica VT1200S, Leica Biosystems, Ontario, Canada). The level of the oblique slice was recorded for each preparation and used in post hoc analysis to determine if SPN calcium imaging responses varied at different thoracic segments. The exposed portion of the spinal cord was then glued to a sylgard-coated (Sylgard, Dow Corning, MI, USA) recording chamber designed and 3D-printed in-house. Specifically, the method is a modification of that described by (Rancic, Haque et al. 2019), in which the obliquely cut surface of the spinal cord was placed on a sylgard “ramp” (∼30°) to enable simultaneous visualization SPNs under fluorescence while maintaining physical continuity with the lumbar spinal cord for generating lumbar locomotor activity, as monitored by ventral root (VR) recordings. All recordings were performed in oxygenated room-temperature aCSF composed of (mM): NaCl (111.0), KCl (3.085), D-glucose (10.99), NaHCO_3_ (25.0), MgSO_4_.7.H_2_O (0.31), CaCl_2_ (2.52), KH_2_PO_4_ (1.1); pH 7.4.

#### Whole cord and lumbar-evoked induction of locomotor activity

In our first series of experiments, whole-bath application of neurochemicals was used to elicit locomotor activity in C57Bl/6 mice (n = 26) with varying combinations of serotonin, NMDA and dopamine, in the following concentrations: 5-hydroxytryptamine (5-HT, Sigma-Aldrich Co., MO, USA; 10–50 µM), N-methyl-D-aspartic acid (NMDA, Sigma-Aldrich; 2–10 µM), and dopamine (DA, Sigma-Aldrich; 50–100 µM). In some experiments di-hydro-kainic acid (DHK, Tocris Bioscience, Bristol, UK; 150–400 µM, n =16 trials in 14 mice), and bicuculline (BIC, Sigma-Aldrich; 10–40 µM, n=4 trials in 4 mice) were also applied to induce rhythmic activity. Neurochemical concentrations refer to final bath concentrations.

Using ChatCre/GCamp6s mice in another series of experiments (n=15), a split-bath preparation was used to isolate the lumbar spinal cord from the thoracic spinal cord with a 3D-printed 2-chamber bath sealed at the chamber barrier and spinal cord contact edges with petroleum jelly (Cowley and Schmidt 1997). A dual-perfusion system was used to apply the locomotor cocktail (20-30 µM 5HT, 5 µM NMDA and 5-10 µM DA) exclusively to the lumbar region, while the thoracic region was perfused with oxygenated recording aCSF. Drugs were then applied only to the lumbar spinal region to evoke locomotor activity which allowed us to record SPN activity in the absence of direct effects of the neurochemicals on SPNs. Red food coloring dye placed in the lumbar chamber following experimentation was used to confirm integrity of seal and that drug application was confined to the caudal bath segment.

#### Lumbar ventral root monitoring and recording

Lumbar VR recordings were obtained using glass suction electrodes (120-140 µm inner diameter). We classified tonic activity as a sustained increase in VR discharge compared to baseline whereas rhythmic locomotor activity was defined by VR bursting between left and right L2 or L5, or between ipsilateral L2 and L5 VRs since L2 corresponds with flexor, and L5 with extensor phases of the step cycle, respectively (Cowley and Schmidt 1994, Kiehn and Kjaerulff 1996). VR signals were band-pass filtered at 30–3000 Hz, amplified (custom-made SCRC amplifier), digitized and acquired in gap-free mode at 2 kHz with a CED Power 1401 AD board, and displayed with Signal 7.6 Software (Cambridge Electronic Devices, Cambridge, UK). All recordings were performed at room temperature (∼22°C). A TTL pulse from the camera to the electrophysiological data capture system (Signal 7.6, Cambridge electronic devices, Cambridge, UK) synchronized calcium imaging with VR recordings.

### Calcium imaging

#### Retrograde labeling of thoracic SPNs

After dissection, spinal cords from neonatal C57BL/6 mice (P0-P5) were transferred (ventral side up) into a chamber with oxygenated room-temperature aCSF. SPNs and somatic MNs from different thoracic segments (T4-T12) were retrogradely labeled by applying calcium dye (Ca-green conjugated dextran amine, CGDA; 3000 MW, Life Technologies Corp., OR USA) to the cut ends of thoracic VRs (Th-VRs, see Fig. 1Ai). CGDA crystals were diluted with aCSF into 20% stock solution (25 µL of aCSF / 5 mg of CGDA crystals). A small volume (4-6µL) of the diluted calcium dye was applied to thoracic VRs using suction glass microelectrodes (100-120µm inner diameter). Thoracic VRs were cut immediately before suctioning to increase dye uptake and sectioned close to their exit from the spinal cord to minimize labelling time. Thoracic VRs were selected based on viability and length. Retrograde labeling of corresponding thoracic MNs and SPNs continued in the dark at room temperature (∼22°C) for at least 3 hours (Szokol, Glover et al. 2008).

**Figure 1.**
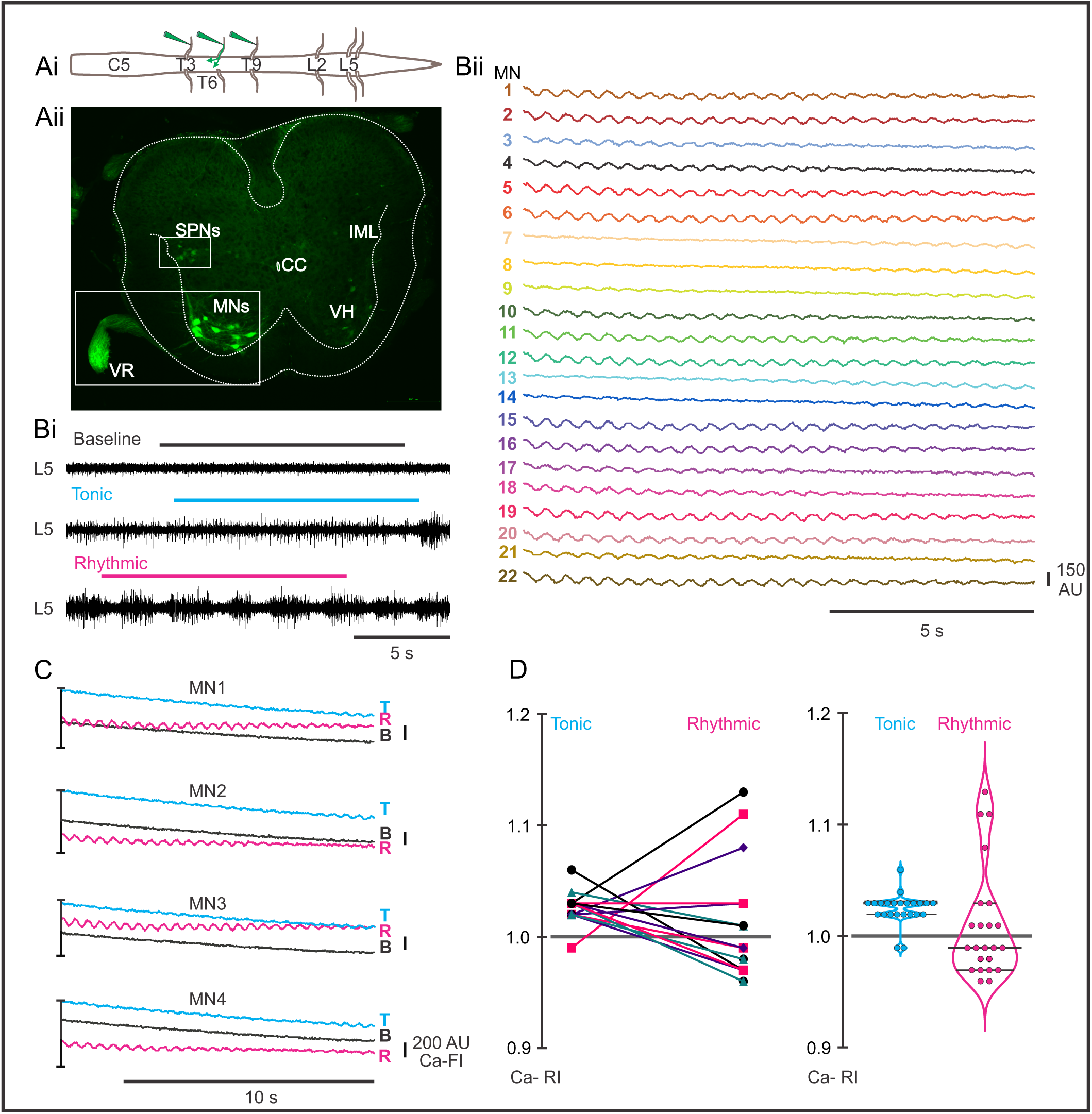
**Calcium responses in thoracic motoneurons (MNs) during tonic and rhythmic ventral root (VR) activity. Ai**. *Schematic* of the retrograde dye labelling procedure and lumbar VRs. **Aii**. Transverse section of thoracic spinal cord with retrogradely labeled MNs and SPNs. **Bi**. VR recordings at baseline, and during tonic and rhythmic activity. **Bii**. calcium imaging responses of 22 motoneurons at T10–11 during rhythmic activity induced by 20 µM 5-HT, 5 µM NMDA, and 50 µM DA. **C.** Ca-FI traces from four representative MNs (MNs 1-4 in Bii), shown during baseline, tonic, and rhythmic VR activity. Ca-FI increased during tonic activity with some transitioning to rhythmic oscillations that became clearly evident during rhythmic activity. Note that absolute intensity levels were higher during tonic versus rhythmic activity for each of these 4 MNs. Calcium intensity is shown in arbitrary units (AU, scale bar bottom right). **D.** Individual Ca-RI (*left panel),* and violin plots (*right panel*) for all 22 MNs during tonic and rhythmic VR activity. Most MNs showed an initial increase in Ca-RI during tonic activity compared to normalized baseline, which may further increase during rhythmic activity. Violin plots summarizing Ca-RI values indicated that while tonic activity led to a general increase in Ca-RI, a subset of MNs exhibited a decrease in Ca-RI during rhythmic activity. Thick black lines within each violin plot represent the median Ca-RI value; thinner black lines indicate the 10th and 90th percentiles. Tonic VR activity median = 1.03, rhythmic VR activity median = 0.99. Note: for this and all other figures, baseline is indicated in black, tonic activity by cyan and rhythmic activity in magenta.

#### Thoracic SPNs visualization

Labeled thoracic SPNs (or MNs), either by retrograde calcium dye or by genetically encoded calcium indicator GCaMP6s, were visualized in obliquely cut cords at different thoracic levels (T3- T12), as described above. The thoracic surface was visualized first using 5x wide objective lens (Axio Examiner.Z1 microscope; Göttingen, Germany) for a boarder view, identifying ventral and dorsal horns and the central canal. Then, under fluorescence with the 20x wide objective, clusters of labeled SPNs were identified based on their lateral location and position relative to the central canal, where the intermediolateral cell column (IML) is located. MNs pools were clearly distinguishable from SPNs based on their size and ventral location in lamina IX (see Fig. 1Aii).

#### Optical Recording of Calcium Responses

Thoracic SPNs were imaged using a 20x wide aperture (1.2 nA) water-immersion objective lens on an upright epi-fluorescence Zeiss Axio Examiner microscope. Fluorescence intensity changes were recorded using Prime BSI Scientific CMOS camera (Photometrics, BC, Canada) and SlideBook 6.0 software (Intelligent Imaging Innovations, Denver, CO, USA, RRID:SCR_014300). Image series were captured at sampling frequencies between 4.5-7.5 Hz, and occasionally at 40 Hz. Regions of interest (ROIs) were drawn around the somas of labeled thoracic SPNs in the IML or MNs in the ventral horn to record fluorescence intensity at baseline (before drug-induced locomotion) and during tonic and/or rhythmic VR activity. We set a background fluorescence region as an area within the imaged field far from labelled thoracic SPNs to normalize for changes in global fluorescence changes over time during each experiment.

### Data analysis

#### Post-hoc normalization and analysis procedures

For all analyses, calcium fluorescence intensities (Ca-FI) for each SPN were exported from Slidebook software into Microsoft Excel. To compare Ca-FI at baseline and during tonic and rhythmic VR-activity for individual SPNs, each CA-FI was normalized to the background fluorescence region in each condition. This allowed us to calculate the relative intensity (Ca-RI) of normalized fluorescence intensity for each ROI during tonic and rhythmic activity and express it as a percentage of baseline fluorescence (Ratliff, Pekala et al. 2023). For some animals, files were converted into text files and imported into either Analysis (in-house SCRC software) or Signal 7.6 Software for subsequent oscillatory calcium event visualization (plotting) and analysis. For MNs, oscillations in Ca-RI within each Slidebook capture window were robust, consistent and easily visually identifiable, particularly when sampling at 40 Hz. Rhythmic events in SPNs were less common and regular, and lower in frequency and intensity, and, as expected, did not correlate directly with rhythmic VR events. Data were exported as text files into GraphPad Prism Software, San Diego, California, USA, for graphing and statistical analysis. Averages are expressed as means ± standard deviations, and ranges denoted in square parentheses [].

## RESULTS

### Ca-RI responses observed in MNs during tonic and rhythmic locomotor activity during whole bath-application of neurochemicals

Initially we recorded activity in 56 CDGA retrogradely labelled thoracic MNs (6 preparations, T7-T10, see dye application and VR recording set up Fig. 1A) and from 19 thoracic MNs that expressed GCaMP6s indicator (4 preparations, T4-5) to test our methodology in comparison to previous studies. As expected, at baseline, CDGA-labelled MNs did not demonstrate oscillatory transients in Ca-FI. Also as expected, we observed regular oscillations in Ca-FI recorded in CDGA-labelled thoracic MNs during rhythmic VR activity (Fig. 1B and C), consistent with others’ findings of rhythmic oscillations in motoneurons and locomotor-related spinal interneurons during *in vitro* locomotor activity (Jean-Xavier and Perreault, 2018; Rancic et al., 2020). During rhythmic VR activity, many of the 22 MNs displayed rhythmic oscillations throughout each 12-s record (e.g., Fig. 1Bii MNs 1-2, 5-6), whereas oscillations in other MNs appeared episodically, occurring at the beginning or end of each record (e.g., Fig. 1Bii, MNs 7-9, 13-14). During tonic VR activity that typically precedes the appearance of rhythmic VR activity, most MNs showed increased Ca-FI, with some also displaying rhythmic oscillations in Ca-FI (e.g., Fig. 1C, cyan). Relative Ca-FI was lower during rhythmic activity in these 4 MNs when compared to tonic discharge (Fig. 1C).

Similarly, most of the 22 MNs examined displayed an increase in Ca-RI during tonic VR activity (Fig. 1D, violin plots, right), likely related to increased resting membrane potentials of MNs observed during tonic VR activity (Krawitz, Fedirchuk et al. 2001, MacDonell, Power et al. 2015, Jean-Xavier and Perreault 2018). During rhythmic VR activity, responses were more variable, with increased Ca-RI in 6/22 (27%), Ca-RI remaining relatively similar to baseline in 9/22 (41%) and decreased Ca-RI in 8/22 (32%) MNs (Fig. 1D, right). The regular MN Ca-RI oscillations observed during rhythmic VR activity likely reflects underlying locomotor drive potentials (LDPs) observed during fictive locomotion (Jordan 1983, Hochman and Schmidt 1998). The MN Ca-RI intensity changes we observed were similar to those observed in thoracic MNs in response to brainstem stimulation (Szokol, Glover et al. 2008), and to those observed in MNs and presumed locomotor-related INs during rhythmic lumbar locomotor activity (O’Donovan, Bonnot et al. 2005, Jean-Xavier and Perreault 2018, Rancic, Ballanyi et al. 2020).

### SPN responses during whole-bath application of neurochemicals

#### Thoracic SPN Ca-RI responses during lumbar VR locomotor activity induced by whole-cord drug application

We used two approaches to characterize thoracic SPNs responses during lumbar locomotor activity. In the first, we perfused the entire spinal cord with neurochemicals to induce fictive locomotion while recording SPN population calcium responses at a variety of rostrocaudal levels (T4 to T11) and monitored SPN activity levels at baseline, and during tonic and rhythmic activity. In our second approach we limited neurochemical application to T13 and caudal to remove the effect(s) of direct neurochemical application on SPNs while maintaining the ability to generate hindlimb locomotor activity (Cazalets, Borde et al. 1995, Kjaerulff and Kiehn 1996, Cowley and Schmidt 1997, Kremer and Lev-Tov 1997, Cowley, Zaporozhets et al. 2009), see below. Calcium image recordings were performed from 313 CGDA labelled thoracic SPNs (25 preparations) and 192 thoracic SPNs in our ChatCre/Gcamp6S mouse (15 preparations) at different thoracic segments in our whole-bath configuration experiments. SPNs displayed a variety of responses at baseline, and during tonic and rhythmic activity. In some preparations, Ca-RI oscillations were observed (e.g., Fig. 2A and B) as well as increases in Ca-RI (Fig. 2C). In this example, all 4 CGDA-labeled SPNs displayed a larger increase in Ca-RI during tonic (cyan) compared to rhythmic (magenta) VR activity, like that seen in MNs (compare Fig. 2B and C to Fig 1C and D). Here and elsewhere, only changes > ± 3% were classified as increased or decreased Ca-RI.

**Figure 2.**
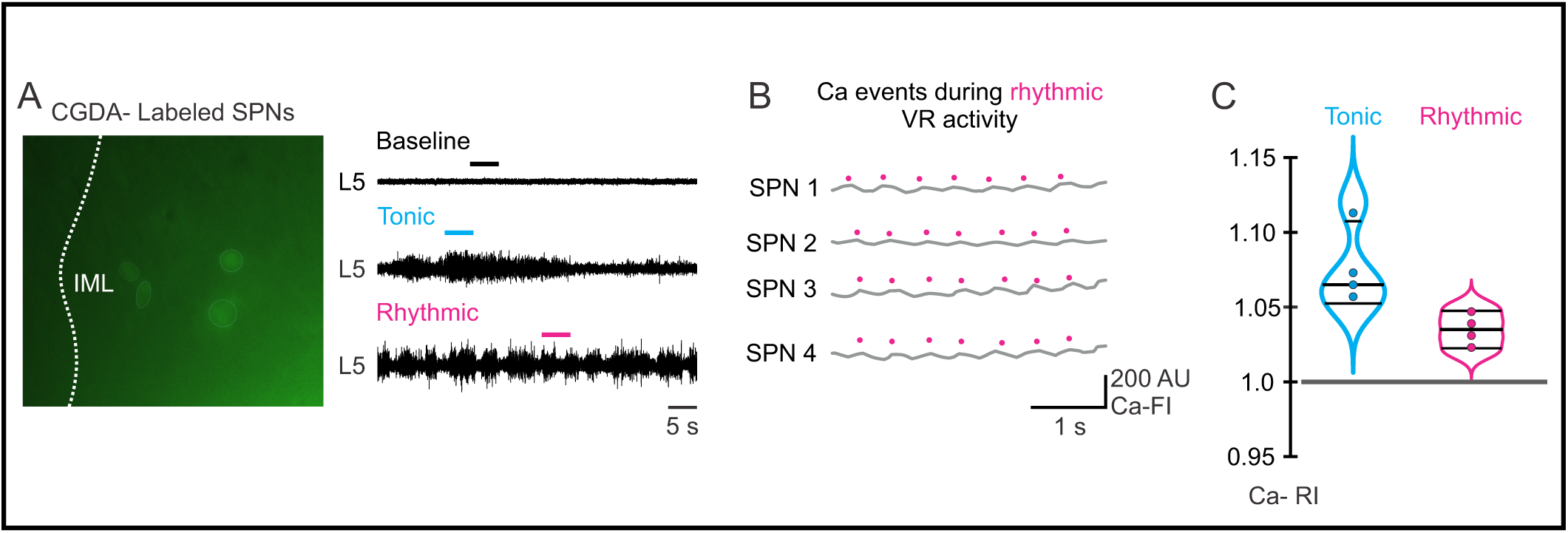
Calcium responses of thoracic sympathetic preganglionic neurons (SPNs) during VR activity induced by whole-bath application of 5-HT, NMDA, and DA. A. *Left panel:* Light circles indicate the ROIs corresponding to SPN cell bodies of four CGDA- retrogradely labeled thoracic SPNs at T6–7. *Right panel:* Corresponding L5 ventral root (VR) recordings and calcium imaging during baseline (black), tonic (cyan), and rhythmic (magenta) activity. **B.** Oscillatory calcium (Ca) events observed in thoracic SPNs during rhythmic VR activity. Magenta dots mark individual Ca events. Calcium intensity is shown in arbitrary units (AU, scale bottom right). Recordings were obtained during a 12 s calcium imaging episode sampled at 7.8 Hz. **C.** Violin plots showing changes in calcium relative intensity (Ca-RI). The gray line at 1.0 indicates the normalized baseline level. Note that Ca-RI increases more during tonic versus rhythmic activity. Thick black lines within each violin plot represent the median Ca-RI value during tonic (median = 1.065) and rhythmic (median 1.035) activity; thinner black lines indicate the 10th and 90th percentiles.

Our focus was to examine changes in Ca-RI at different thoracic segmental levels during tonic and rhythmic lumbar locomotor activity. In the preparation shown in Fig. 3, recordings were obtained at T4/5 before and after whole-bath application of 20µM 5HT and 5µM NMDA (schematic of set up in Fig. 3A). Some Ca-RI increases were readily evident upon visual inspection of the calcium images (e.g., Fig. 3B and C) whereas others required normalization to averaged baseline values and analyses to illustrate changes in Ca-RI (e.g., Fig. 3D). The inserts in Fig. 3Ci-Ciii show that most of the visualized thoracic SPNs (yellow squares in Fig. 3Bi-Bii) progressively increased their Ca-FI during tonic and then rhythmic activity. Two SPNs are highlighted (Figure 3Ci-iii, circled). Overall, most SPNs in this preparation displayed increased Ca-RI during tonic and rhythmic activity, with only one SPN showing decreased Ca-RI compared to baseline during tonic (purple line/dot) and another SPN with decreased Ca-RI compared to baseline during rhythmic activity (teal line/dot; Fig. 3D left).

**Figure 3.**
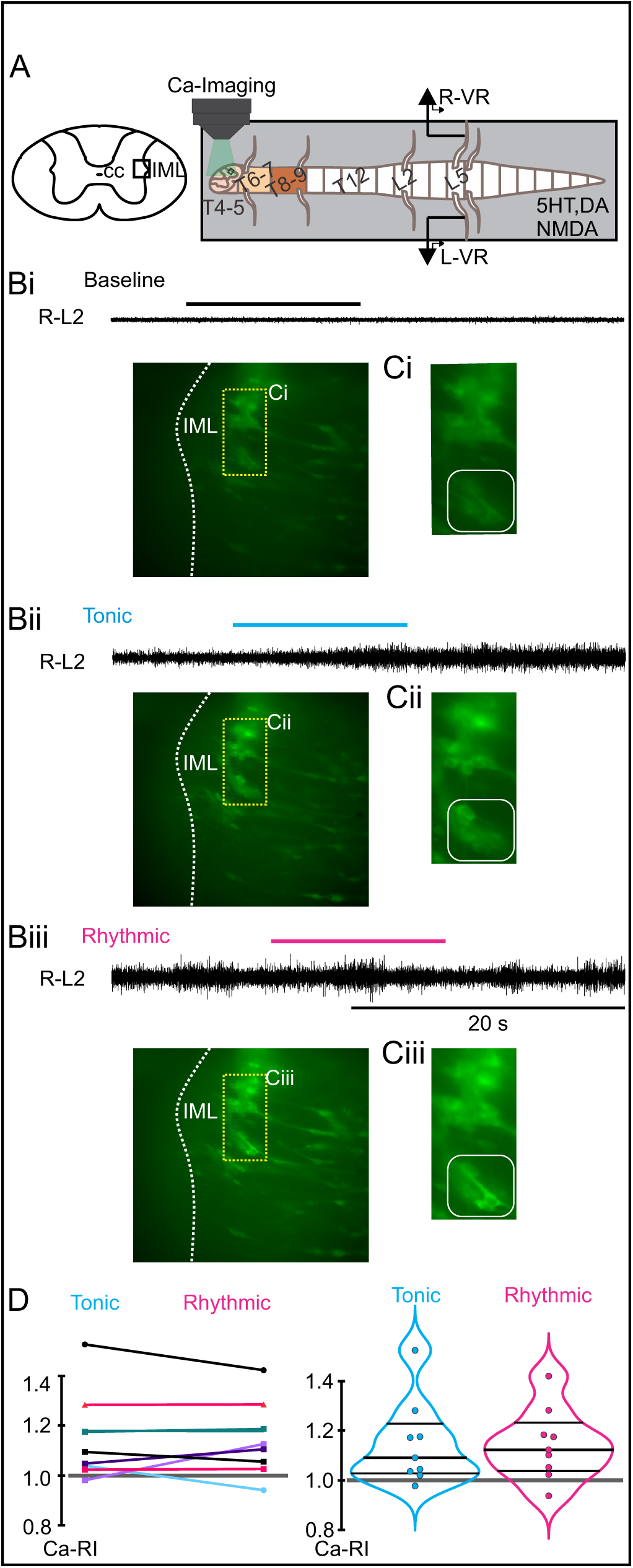
Increasing recruitment of thoracic SPNs during tonic and rhythmic VR activity. **A.** *Left panel:* Schematic of an oblique thoracic spinal cord section showing the IML where SPNs were visualized. *Right panel:* Experimental design for whole bath configuration. Oblique sections were made to expose SPNs at various thoracic levels: T4–5 (peach), T6–7 (yellow), and T8–9 (brown). A locomotor cocktail (5-HT, DA, and NMDA) was applied to the whole spinal cord. Lumbar VR activity was recorded to monitor tonic and/or rhythmic activity, while calcium imaging was simultaneously performed on thoracic SPNs. **B.** Representative calcium imaging of SPNs at the T4–5 level in a GCaMP6s P2 mouse and VR recordings during baseline (**Bi**), tonic (**Bii**), and rhythmic (**Biii**) activity following whole-bath application of 20 µM 5-HT and 5 µM NMDA. **C.** Magnified view of a cluster of T4–5 SPNs at baseline (**Ci**), during tonic (**Cii**), and rhythmic VR activity (**Ciii**). As highlighted by the white square, these two SPNs exhibit a progressive increase in calcium fluorescence during tonic and rhythmic activity. **D.** *Left panel:* Individual Ca-RI for all 9 SPNs during tonic and rhythmic VR activity. Most SPNs show an increase in Ca-RI during both tonic and rhythmic VR activity. *Right panel:* Violin plots summarizing Ca-RI values for the SPNs population during tonic and rhythmic magenta activity. In this example SPNs show a similar increase in Ca-RI during both tonic and rhythmic VR activity. Thick black lines represent the median Ca-RI value; thinner black lines indicate the 10th and 90th percentiles. Tonic VR activity median = 1.09, rhythmic VR activity median = 1.10.

#### SPNs exhibit distinct patterns of Ca-RI responses at different thoracic segmental levels and differed during tonic versus rhythmic lumbar locomotor activity

We observed differences in the pattern of Ca-RI responses during tonic and rhythmic VR activity at different spinal segments (Fig 4A-B). For example, during whole bath drug application, most SPNs recorded from the T4-5 segments (Fig. 4C, left) exhibited increased Ca-RI above the normalized baseline value during tonic VR activity (top) whereas SPN Ca-RI changes during rhythmic activity were mixed (magenta violin plot in Fig. 4C, left). Ca-RI responses at T6-7 were generally increased during rhythmic activity compared to tonic activity (e.g., Fig. 4C middle). SPN recordings from more caudal thoracic segments (i.e., T8 through T11) showed mixed responses with half showing increased and half showing decreased Ca-RI during both tonic and rhythmic VR activity (Fig. 4C right).

**Figure 4.**
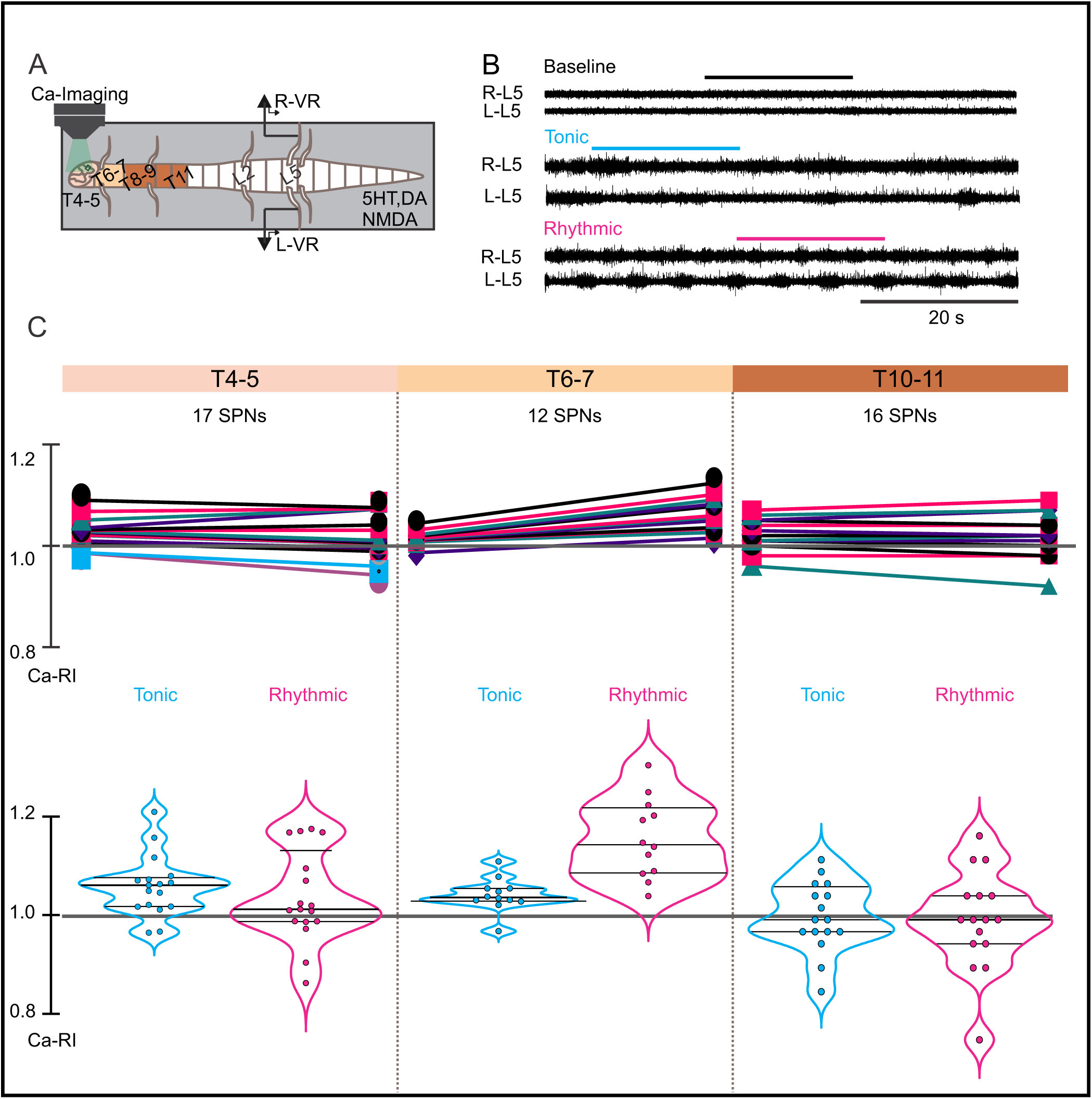
Thoracic SPN Ca-RI changes at different segmental levels after whole-cord application of locomotor cocktail. **A.** Experimental set up for whole cord configuration, similar to Figure 3. **B.** Calcium imaging and VR recordings during baseline, tonic and rhythmic VR activity at T6-7. **C**.Ca-RI changes for 3 different experiments recorded at each thoracic level color-coded by spinal segment (T4-5, T6-7, T10-11). Upper panels represent the individual Ca-RI changes during tonic and rhythmic activity. Most of the SPNs at T4-5 show an increased Ca-RI during tonic and rhythmic activity. SPNs at T6-7 displayed a larger increased Ca-RI during rhythmic compared to tonic activity and most SPNs at T10-11 no change in Ca-RI. Violin plots show Ca-RI values distribution as population in each SPN cluster by thoracic segment. Thick black lines within each violin plot represent the median Ca-RI value; thinner black lines indicate the 10th and 90th percentiles. While a slight increase in Ca-RI is observed during tonic activity for SPNs at the T4-5 level, (median = 1.031), SPN responses during rhythmic activity were more variable, with 5 increasing, 2 decreasing, and most remaining similar to baseline (median = 1.01). In contrast, most SPNs recorded at T6-7 show a greater increase in Ca-RI during rhythmic compared to tonic activity. Tonic VR activity median = 1.012 and median = 1.057 during rhythmic VR activity. SPNs at T10-11 displayed Ca-RI values around baseline (median = 1.02) during tonic activity and rhythmic activity (median 1.01)

In summary, we recorded 164 SPNs at the T4-5 level (11 preparations, 12 trials) during tonic VR activity induced by whole cord drug application, and Ca-RI increased in 82/164 (50%), was unchanged in 57/164 (35%), and decreased in 25/164 (15%). Of the 153 thoracic SPNs recorded at T4- 5 during rhythmic VR activity (9 preparation, 12 trials), Ca-RI increased in 51/153 (33%), was unchanged in 53/153 (35%), and decreased in 49/153 (32%). Thus, while 50% of SPNs in T4-5 segments showed increased Ca-RI during tonic VR activity, fewer (33%) increased Ca-RI during rhythmic VR activity.

We recorded 47 SPNs at the T6-7 level during tonic VR activity (5 preparation, 6 trials), and Ca-RI increased in 23/47 (49%), was unchanged in 23/47 (49%), and decreased in 1/47 (2%). Of the 39 thoracic SPNs recorded at T6-7 during rhythmic VR activity, Ca-RI increased in the majority of SPNs 87%, 34/39) and was unchanged in 5/39 (13%). Thus, a greater proportion of SPNs at T6-7 displayed increased Ca-RI during rhythmic (87%) compared to during tonic (49%) activity with whole bath drug application.

Only a small proportion of recorded SPNs displayed increased Ca-RI at caudal thoracic levels. During tonic activity, only 7% of SPNs (6/82) had increased Ca-RI at T8-9 levels (8 preparations, 9 trials) and 17% (9/114) at T10-11 (12 preparations, 14 trials). Similarly, low numbers of SPNs displayed increased Ca-RI during rhythmic activity at these levels. Specifically, 11/93 (12%) SPNs at

T8-9 and 20/167 (12 %) at T10-11 levels displayed increased Ca-RI during rhythmic activity. Most commonly, SPNs at T8-T11 showed no change in Ca-RI during either tonic or rhythmic activity (77% at T8-9 and 74% at T10-11).

### SPN responses during split-bath application of neurochemicals

#### Ca-RI responses in thoracic SPNs during tonic and rhythmic locomotor activity induced by selective application of neurochemicals to the lumbar spinal cord

To determine whether direct effect(s) of neurochemicals on SPNs and thoracic spinal neural circuitry contributed to SPN Ca-RI responses during tonic and rhythmic VR activity, in our second approach, we removed the potential direct effects of the neurochemicals on SPNs by using a split-bath configuration, in which locomotor-inducing neurochemicals were applied selectively below the T12/13 spinal level. Thus, the split-bath configuration enabled perfusion of neurochemicals to the chamber containing the lumbosacral spinal cord to activate only the lumbar portion of the locomotor CPG, thereby enabling assessment of the influence of ascending propriospinal projections to thoracic SPNs during lumbar-evoked locomotor activity. We recorded calcium responses from 191 thoracic SPNs (15 preparations) from different thoracic segments at baseline and during tonic and rhythmic VR activity (Fig. 5A-B).

**Figure 5.**
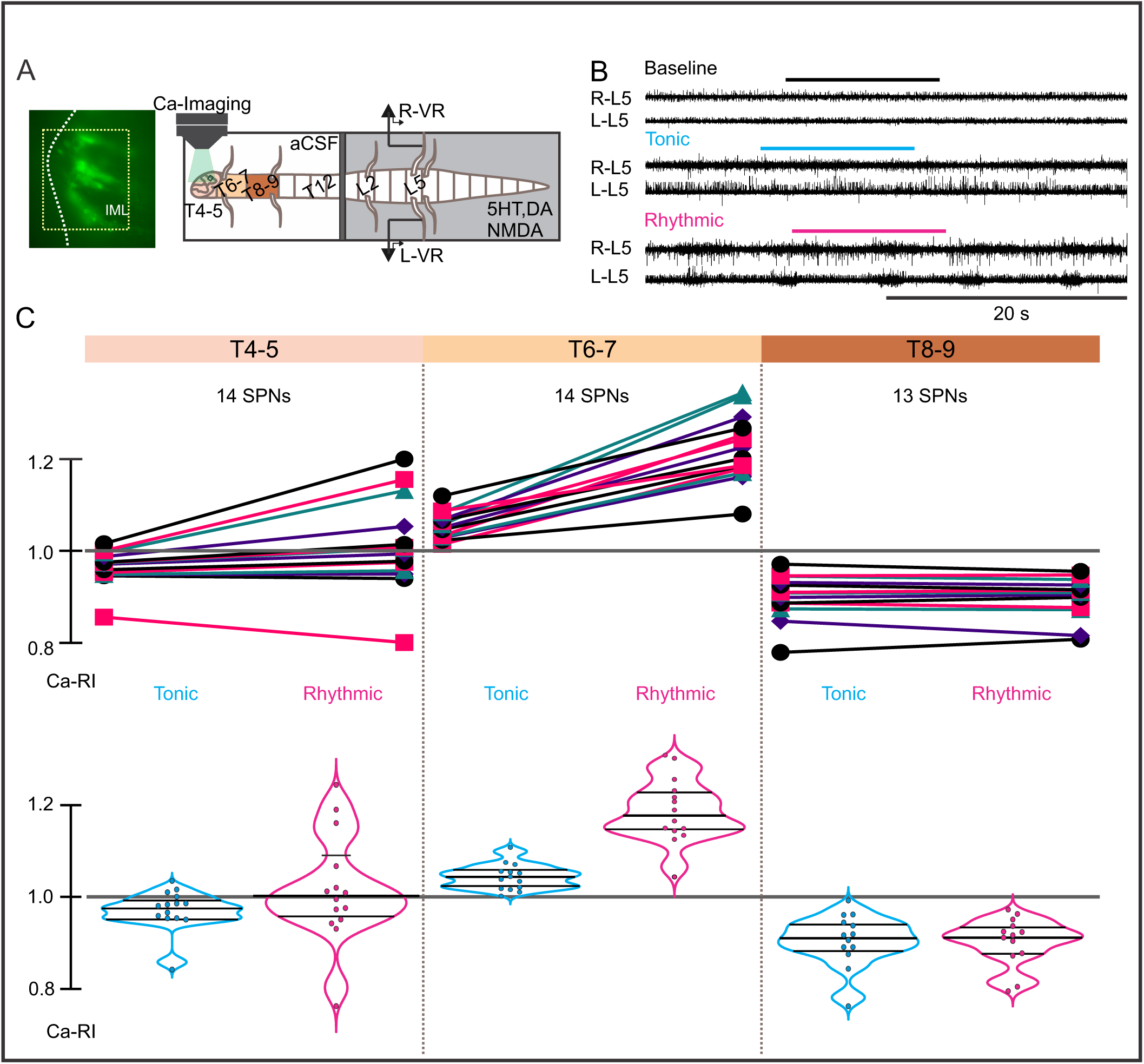
Ca-RI changes in thoracic SPNs during tonic and rhythmic versus VR activity induced by lumbar application of neurochemicals by thoracic level. **A**. Experimental set up. Left panel. Calcium imaging of thoracic SPNs at T6-7 level in a GCaMP6s P4 mouse during rhythmic VR activity. Right panel. Split-bath configuration for selective pharmacological activation of lumbar spinal neurons with barrier placed between T13 and L1 spinal cord segments to physically separate thoracic and lumbar regions (gray rectangle). **B.** Calcium imaging at T6-7 and L5 VR recordings during baseline, tonic and rhythmic VR activity (same preparation as in A right panel and C middle panel). C.Ca-RI changes for 3 different experiments recorded at each thoracic level color-coded by spinal segment (T4-5, T6-7, T8-9). Upper panels represent the individual Ca-RI changes. Although some SPNs at T4-5 level (left panel) show a slight increased Ca-RI during tonic activity and greater intensity changes are observed during rhythmic activity. As shown in this particular example, SPNs at T6-7 segment (middle panel), exhibited a larger increase in Ca-RI during rhythmic versus tonic VR-activity. However, SPNs at T8-9 segment displayed decreased Ca-RIs during both tonic and rhythmic VR activity. Lower panels. Violin plots show Ca-RI values distribution as population in each SPN cluster by thoracic segment. Thick black lines within each violin plot represent the median Ca-RI value; thinner black lines indicate the 10th and 90th percentiles. T4-5 segment median value = 0.95 during tonic, and median = 1.01 during rhythmic VR activity. T6-7 segment median value = 1.059 during tonic, and median = 1.22 during rhythmic VR activity. T8-9 segment median value = 0.90 during tonic, and median = 0.89 during rhythmic VR activity.

Thoracic SPNs also exhibited divergent Ca-RI responses during lumbar-induced tonic and rhythmic VR activity at different spinal levels (Fig. 5C). These calcium responses showed some similar trends to those observed during whole-bath application of locomotor-inducing neurochemicals in that SPN Ca-RI was increased at T6-7 levels during rhythmic activity (Fig. 5C, middle), whereas SPN Ca- RI was mainly decreased at caudal thoracic levels (T8-9, Fig. 5C right) during tonic and rhythmic activity. Note that in the absence of direct neurochemical activation, in contrast to that seen with whole bath application, SPN Ca-RI at T4-5 was either decreased or unchanged during tonic activity. SPN responses at T4-5 were similar during rhythmic activity in the split and whole bath configurations, such that similar proportions of SPNs showed increased, unchanged or decreased Ca-RI (compare Fig. 5C, left with Fig. 4C, left).

#### SPNs exhibit distinct patterns of Ca-RI responses at different thoracic segmental levels that differed during tonic versus rhythmic lumbar locomotor activity

In summary, in the split-bath configuration, at T4-5, we recorded Ca-RI in 36 SPNs during tonic and in 59 SPNs during rhythmic VR activity. During tonic activity at T4-5, Ca-RI in SPNs was either unchanged (47%, 17/36) or decreased (53%, 19/36). During rhythmic activity at T4-5, Ca-RI was increased in a minority of SPNs (12%, 7/59) whereas the majority were either unchanged (24%, 14/59 or had decreased Ca-RI (64%, 38/59). We recorded 65 SPNs at T6-7 during tonic VR activity, and approximately one-third (34%, 22/65) showed increased Ca-RI, most remained unchanged (48%, 31/65), and a small portion of SPNs had decreased Ca-RI (18%, 12/65). We recorded Ca-RI from 76 T6-7 SPNs during rhythmic VR activity and Ca-RI was increased in the majority of SPNs 53/76 (70%), and was either unchanged (13/76, 17%) or decreased (10/76, 13%) in the remainder of SPNs. At T8- 9, Ca-RI was increased in only 2/30 (7%) of SPNs during tonic activity, remaining unchanged (30%, 9/30) or decreased (63%, 19/30) in the majority of SPNs. During rhythmic VR activity 100% of recorded SPNs at T8-9 (25/25) showed decreased Ca-RI.

#### Comparison of rostrocaudal spatial distribution of Ca-RI responses in thoracic SPNs during whole- and split-bath application of neurochemicals

Results from the whole and split-bath series of experiments are summarized in Figure 6. Overall trends in the SPN responses at rostral versus caudal thoracic spinal levels were similar between the whole-bath and split-bath series of experiments. As shown, the greatest proportion of thoracic SPNs exhibiting an increase in Ca-RI during either tonic or rhythmic lumbar VR activity were located at the T6-7 level, which is particularly evident from the split-bath series of experiments (Fig. 6Bi and Bii). During rhythmic activity (Fig. 6Aii *vs* Bii), although a greater proportion of SPNs at T6-7 had increased Ca-RI in the whole cord versus split bath configuration (87%, 34/39 in whole vs 70%, 53/76 in split; Fig. 6Aii *vs* Bii), the overall averaged mean change in Ca-RI was greater at T6-7 in the split-bath configuration (1.22 ± 0.11 [1.03-1.42] in lumbar-only *versus* 1.13 ± 0.06 [1.04-1.27] in whole cord application).

**Figure 6.**
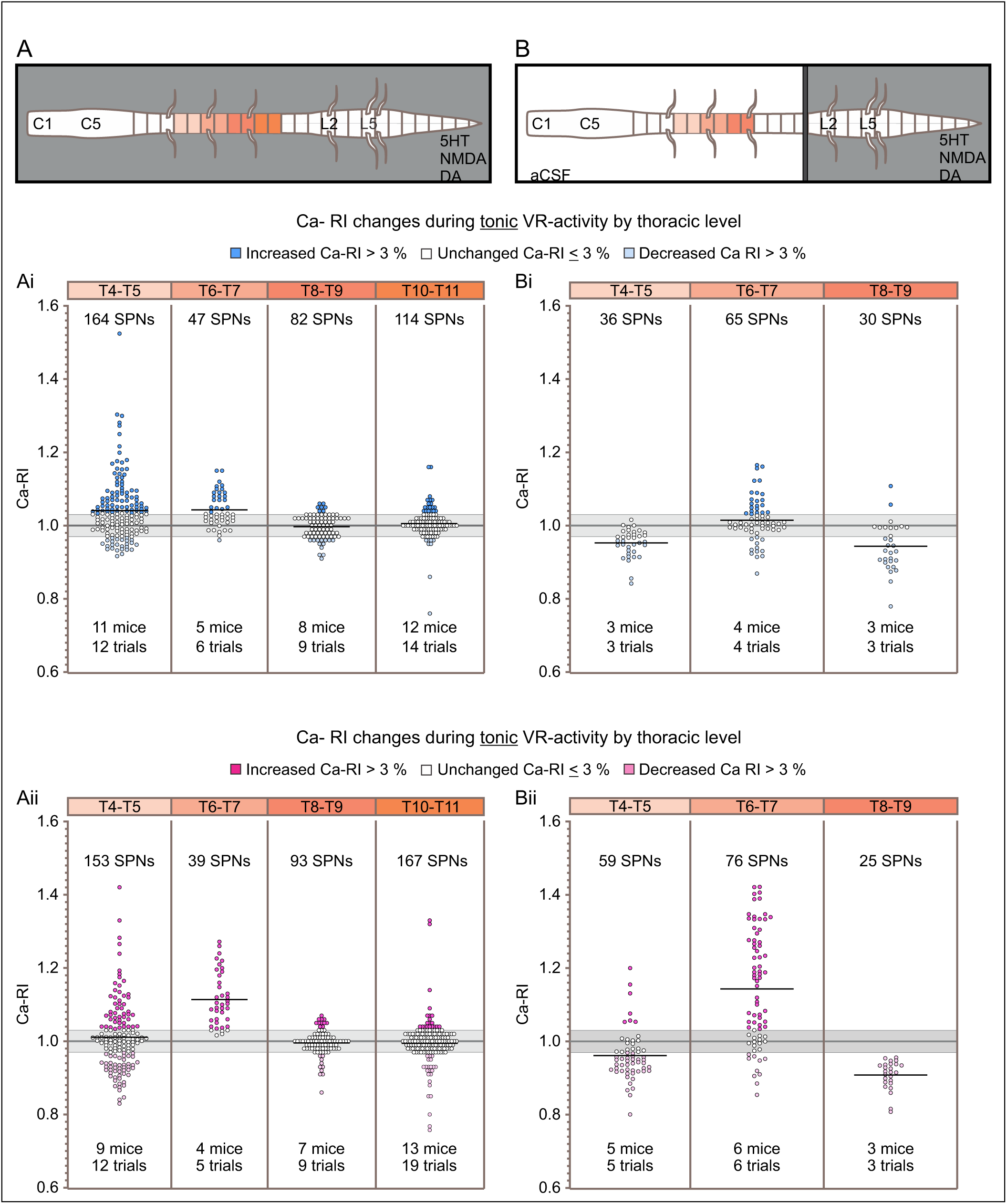
Thoracic SPNs show a differential rostro-caudal distribution of Ca-RI depending on whether 5-HT, NMDA, and DA are applied to the whole spinal cord or only to the lumbar/sacral region. Schematics of the bath design for whole-cord (**A**) and lumbar-only (**B**) application of 5-HT, NMDA, and DA to induce tonic (cyan) and/or rhythmic (magenta) VR activity. Ca-RI changes are summarized for all thoracic SPNs recorded at each segmental level and color-coded by segment. For each panel in Ai through Bii, the gray line represents the normalized baseline (1.0), and the gray shadowed region indicates RI changes < or > 3%, and SPNs in this range are white. During tonic VR activity (**Ai, Bi**), SPNs showing increases >3% are coloured dark blue, and decreases >3% as light blue. During rhythmic VR activity (**Aii, Bii**), SPNs showing increases >3% are coloured dark pink, and decreases >3% as light pink. In the whole bath configuration for both tonic and rhythmic VR activity, greater numbers of SPNs showed increased Ca-RI responses at rostral thoracic levels (T4 – T7) versus caudal levels (T8-11) (**Ai and Aii**). Note the greater increases in SPN intensity at T6-T7 during rhythmic activity (compare Aii to Ai). Similar trends were observed in SPNs with lumbar-only application of neurochemicals such that greater numbers of SPNs in rostral segments showed increased Ca-RI during both tonic and rhythmic VR activity compared to caudal thoracic segments (**Bi and Bii**). However, note that during lumbar-only application, the greatest numbers of SPNs with increased Ca-RI centered upon T6-7.

Examining SPN responses at different segmental levels *in the split-bath configuration* indicated that activity in the majority of SPNs at segmental levels other than T6-7 either remained unchanged or decreased during tonic or rhythmic activity. Specifically, at T4-5 during tonic activity, SPNs had either decreased Ca-RI (53%, 19/36) or unchanged Ca-RI (47%, 17/36), with overall averaged mean change of 0.93 (± 0.03 [0.84 - 0.96]). At T4-5 during rhythmic activity, the majority of SPNs had either decreased Ca-RI (64%, 38/59) or unchanged Ca-RI (24%, 14/59) with only 12% of SPNs with increased Ca-RI (overall mean = 0.92 ± 0.03 [0.80 - 0.97]). At T8-9 during tonic activity, the vast majority of SPNs had either decreased 63% (19/30) or unchanged 30% (9/30) Ca-RI, with only 2/30 (7%) showing increased Ca-RI (overall mean = 0.91 ± 0.04 [0.81-0.96]). Finally, during rhythmic activity at T8-9, 100% of SPNs (25/25) had decreased Ca-RI (0.91 ± 0.04 [0.81- 0.96]) during locomotor activity.

Except for SPN responses at the T6-7 segmental level, whole cord application of neurochemicals led to greater numbers of SPNs showing increased Ca-RI at every level examined, during either tonic or rhythmic activity, when compared to lumbar-only drug application. These observations could be explained by direct effect(s) of the neurotransmitter receptor agonists used to elicit locomotor-like activity on thoracic SPNs, since each of dopamine, serotonin and NMDA have been reported to depolarize SPNs in slice *in vitro* (Spanswick and Logan 1990, Pickering, Spanswick et al. 1994, Gladwell and Coote 1999, Zimmerman, Sawchuk et al. 2012). In viewing SPN responses for the whole-bath series during tonic VR activity it is apparent that a greater proportion of recorded SPNs had increased Ca-RI at all spinal levels when compared to the split-bath series (compare Fig. 6Ai to 6Bi), likely due to direct effect(s) of these neurochemicals on SPN excitability in addition to any changes caused by propriospinal activation during lumbar locomotor network activity. In further support of this, in the split bath configuration, Ca-RI in the vast majority of SPNs at either T4-5 or T8- 9 either remained unchanged or decreased *during tonic activity* (Fig. 6 Bi; 100% at T4-5 and 93% at T8-9). Similarly, Ca-RI in the majority of SPNs at either T4-5 or T8-9 remained unchanged or decreased during *rhythmic VR activity* (88% at T4-5; 100% at T8-9%).

Chi-square analysis indicated that the likelihood of these response patterns being due to chance for each of the whole-bath and split-bath configurations, during either tonic or rhythmic activity was extremely low, p < 0.0001 (Contingency tables can be seen in supplemental data).

## DISCUSSION

The goal of this study was to examine whether activity within lumbar spinal locomotor networks can increase activity in thoracic SPNs. We previously demonstrated a direct ascending excitatory connection from a class of lumbar locomotor-related interneurons (V3 INs) to thoracic SPNs. We hypothesized that, similar to the integration seen at the brainstem level (Koba, Kumada et al. 2022), spinal neurons are also capable of integrating locomotor network activity with sympathetic networks (SPNs), likely mediated from ascending lumbar propriospinal neurons. To achieve this, we recorded lumbar ventral root activity and calcium events from thoracic SPNs in an *in vitro* spinal cord preparation, using either a whole-bath or split-bath configuration. This preparation also allowed us to characterize SPN responses during locomotor activity in the absence of descending commands and afferent input. SPN responses were characterized by rostrocaudal segmental level, in either the presence of locomotor-inducing neurochemicals applied to the entire spinal tissue, or when applied only to the lumbar spinal region. We observed a distinctive pattern in SPN responses at different thoracic segmental levels, with mixed responses in rostral thoracic segments, increased excitability in SPNs at T6-7 and decreased excitability in SPNs at caudal segmental levels.

### Neurotransmitter agonists elicit distinct responses in SPNs, at different thoracic segments

Known rostrocaudal differences in receptors and axon terminal projections may contribute to the differences we observed in SPN excitability at different thoracic levels. NMDA, 5HT and DA have all been reported to depolarize SPNs either directly or indirectly (Spanswick and Logan 1990, Pickering, Spanswick et al. 1994, Gladwell and Coote 1999, Zimmerman, Sawchuk et al. 2012). In *in vitro* studies, glutamate activates SPNs through NMDA and non-NMDA receptors (Spanswick and Logan 1990). Using retrograde labelling and ultrastructural analysis aimed at comparing the proportion of GABA versus glutamatergic synaptic input to SPNs, Llewellyn-Smith and colleagues observed that glutamatergic neurons provide approximately 2/3 of input to SPNs projecting to adrenal glands (T6- T10) and about ½ of input to SPNs projecting to the superior cervical ganglion (T1 – T3) which provides sympathetic input to the head, neck and heart (Strack, Sawyer et al. 1988, Llewellyn-Smith, Phend et al. 1992, Llewellyn-Smith, Arnolda et al. 1998). Llewellyn-Smith and colleagues also showed that virtually all SPNs projecting to either the superior cervical ganglion or the adrenal medulla received input from neuron terminals containing either glutamate or GABA (Llewellyn-Smith, Arnolda et al. 1998). Other sources of synaptic input to SPNs were not examined in these studies (e.g., 5HT input).

Whereas activation of SPNs with NMDA is simply excitatory, the effects of DA and 5-HT are more complex. In response to bath-applied DA, SPNs in mid to upper thoracic spinal cord respond with either a slow hyperpolarization (at ∼95s, 46%), a slow depolarization (at ∼65s, 28%) or a biphasic response (slow hyperpolarization followed by a depolarization, or vice versa, ∼33%) (Gladwell and Coote 1999). The hyperpolarization and depolarization in response to DA was blocked by D1 and D2 antagonists, respectively. (Gladwell and Coote 1999). In caudal segments (T8-L2), DA generally increased excitability in recorded SPNs (5/7) but hyperpolarization and decreased excitability was also noted (2 of 7 SPNs) (Zimmerman, Sawchuk et al. 2012). DA-containing terminals originating from the ipsilateral A11 nucleus are seen at all spinal levels, with particularly dense projections at rostral and caudal thoracic levels, terminating in the IML, around the central canal and in lamina VII (Skagerberg and Lindvall 1985, Yoshida and Tanaka 1988). Thus, these distinct patterns in DA input to thoracic SPNs support our observations of distinct SPN Ca-RI responses at different thoracic segmental levels. The effect of exogenous 5-HT application on SPNs is generally increased excitability characterized by slow and long lasting depolarizations or increased firing rates in >90% of SPNs in slice, and it can induce rhythmic oscillations, or increase the amplitude of spontaneous oscillations (without reported effects on firing rates) in SPNs from whole cord preparations (Pickering, Spanswick et al. 1994, Pierce, Deuchars et al. 2010, Zimmerman, Sawchuk et al. 2012). Variations in the density of descending 5-HT projections at different segmental levels within thoracic IML has been observed in multiple species (as reviewed by (Jensen, Llewellyn-Smith et al. 1995)). In rat for example, the sympathetic nuclei in rostral and caudal thoracic regions contain more 5-HT fibers than mid-thoracic segments (T6 – T10) (Newton and Hamill 1988, Newton and Hamill 1989). Similarly, segmental variations in the density of 5-HT projections and responses also vary within locomotor neural circuits, as evidenced by distinct patterns of segmental expression of serotonergic receptors and distinct functional responses at different rostrocaudal locations [reviewed in (Schmidt and Jordan 2000) and see (Liu and Jordan 2005)]. Thus, SPNs and related autonomic neurons within the IML are directly sensitive to exogenous NMDA, 5HT and DA application, and the responses seen during whole bath application may vary, depending upon rostro-caudal segmental level, receptor subtypes expressed on distinct populations of SPNs, in addition to any ongoing functional activity within locomotor or other neural circuitries. For example, we noted that during tonic activity with whole bath application of drugs, a greater proportion of SPNs had increased Ca-RI, and overall mean changes in Ca-RI were increased in when compared to split bath (lumbar restricted) drug application (compare mean changes in Ca-Ri for each level examined T4-5, T6-7 and T8-9 in Fig. 6Ai *vs* Bi). This suggests that a direct effect on SPNs contributed to the increased Ca-RI seen during tonic VR activity with whole-cord drug application.

In addition to rostrocaudal segmental distinctions, responses in individual SPNs within a single segment were seen, which may be related to distinct tissue or organ targets of each SPN, or to within target distinctions in SPN projections. As discussed by Llewellyn-Smith, there is a link between the neurochemical profiles of inputs to and within SPNs and their functions at the level of the target organ or tissue. This includes for SPNs projecting to the same organ, such that distinct subpopulations with different functions receive distinct combinations of inputs and synthesize and release unique combinations of neuropeptides and other neurotransmitters in addition to acetylcholine (Llewellyn-Smith 2009). As noted by Llewellyn-Smith et al., 1998, that there is clear evidence that neural pathways that control descending sympathetic outflow are organized in the medulla and spinal cord on the basis of function (Llewellyn-Smith, Arnolda et al. 1998). Thus, it would seem reasonable that the pattern of ascending input to SPNs during lumbar locomotor activity would be generally reinforcing or follow the pattern of exciting/increasing activity in SPNs projecting to tissues and organs involved in the ‘fight or flight’ arm of the autonomic nervous system and would simultaneously inhibit or reduce activity in the ‘rest and digest’ arm.

### Ascending Intraspinal Mechanisms for Integration within and between Locomotor and Autonomic Systems

Previous studies have shown that lumbar V3 interneurons exhibit both ascending and descending commissural projections during embryonic development (Blacklaws, Deska-Gauthier et al. 2015, Deska-Gauthier, Borowska-Fielding et al. 2020). Although most V3 INs eventually become local commissural neurons, some evidence suggests that a subset located in the deep dorsal horn extends long ascending projections to cervical CPGs (Zhang, Shevtsova et al. 2022). Moreover, lumbar spinal circuits have been shown to modulate the activity of reticulospinal neurons during locomotion from birth, indicating the presence of ascending projections from lumbar locomotor CPGs (Oueghlani, Simonnet et al. 2018). Supporting this, studies in neonatal rodent preparations demonstrated that ascending locomotor pathways can evoke locomotor-like activity in more rostral spinal segments (Juvin, Simmers et al. 2005, Cherniak, Etlin et al. 2014, Anglister, Cherniak et al. 2017). In contrast, cervical CPGs are less effective at generating rhythmic activity in lumbar segments (Cowley, Zaporozhets et al. 2008, Juvin, Le Gal et al. 2012). Notably, Morin and colleagues proposed that both excitatory and inhibitory ascending propriospinal pathways are required for proper interlimb coordination during locomotion, based on their findings that excitatory input alone was insufficient to produce alternating rhythmic patterns (Juvin, Simmers et al. 2005). Similarly, we observed increased Ca-RI at T6-T7 and decreased Ca-RI at T4-5 and T8-9 SPNs, which suggests that both ascending excitatory and inhibitory components may also contribute to regulation of sympathetic output during lumbar locomotor activity in a segment-specific manner. Thus, it is also possible that rhythm generation depends upon propriospinal connections between locomotor-generating and sympathetic output neurons. In future it would be of interest to determine if locomotor rhythm generation is dependent upon feedback from spinal sympathetic circuitry.

### Limitations and other considerations

In our restricted drug application to the lumbar spinal cord region we would also have directly activated SPNs in the most caudal part of the sympathetic network (i.e., in L1 and L2). Given that gap junctions exist between SPNs throughout the spinal sympathetic IML, although unlikely, it is possible that our drug application may have influenced Ca-RI in the rostral bath via gap junction-mediated communication between SPNs alone, rather than because of activating lumbar locomotor circuitry. However, if our observations were due to directly activating SPNs within these caudal sympathetic nuclei, and then mediated via gap junction signaling to more rostral regions, we would expect to see the same pattern of responses during whole and split-bath drug application, and we did not. Additionally, although gap junction-mediated communication may play a role in synchronizing activity in SPN networks, we previously demonstrated that at least one class of locomotor-related lumbar spinal neurons (i.e., V3 neurons) provide direct synaptic excitatory input to SPNs, indicating the presence of classical synaptic communication elements between locomotor and sympathetic neural circuits in the spinal cord (Chacon, Nwachukwu et al. 2023). Future experiments recording from SPNs directly, in split-bath preparations and thoracic application of gap junction blockers could be used to determine relative contributions of gap junctions to the responses reported here.

### Conclusions

These findings indicate the presence of functional intraspinal communication elements between spinal locomotor and sympathetic systems. Such coordinating communication likely contributes to increased activation of supportive homeostatic and metabolic body tissues and organs (e.g., increased BP or lipolysis). For example, a potential role(s) for this communication pathway may be to provide an ascending spinal component that contributes to the ‘exercise pressor reflex’ in which cutaneous and muscle afferents of the lower limb increase respiratory and cardiovascular activity (Coote, Hilton et al. 1971, McCloskey and Mitchell 1972). We also observed decreases in activation of SPNs in caudal thoracic segments, which may relate to decreased sympathetic drive to tissues and organs whose activity is normally decreased during movement and exercise (e.g., digestion, urine production). Further research would determine whether the decreased excitation of SPNs we observed in more caudal thoracic SPNs during lumbar locomotor activity contributes to these decreases. Ascending intraspinal communication between lumbar locomotor and thoracic sympathetic circuitry may also serve to reinforce integration between ongoing movement and activation of related homeostatic and metabolic sympathetic target tissues and organs normally activated by descending central autonomic commands during movement and exercise. Together, this coordinated ascending intraspinal communication could serve to assist in the excitation and inhibition of body functions needed to sustain ongoing movement at a range of intensities and durations.

Our split-bath results suggest SPN calcium responses can be evoked through activation of lumbar CPGs, indicating that spinal locomotor networks retain the ability to modulate SPN activity via ascending propriospinal connections. While descending motor and autonomic control is dominant in intact systems, our findings suggest that ascending propriospinal input contributes to modulating SPN activity during locomotion, particularly in the absence of supraspinal or sensory input. Hence, we propose that locomotor-sympathetic intraspinal pathways contribute to an integration system that regulates sympathetic outflow during locomotion and functions as a compensatory feedback mechanism to help provide homeostatic support during locomotion. Such mechanisms may become therapeutically useful after spinal cord injury, where restoring or controlling below-injury autonomic functions remain a major challenge.

## List of Abbreviations

5-HT: Serotonin
aCSF: Artificial cerebrospinal fluid
Ca-FI: Calcium fluorescence intensity
Ca-RI: Calcium relative intensity
CDGA: Calcium dextran green amine
CPG: Central pattern generator
DA: Dopamine
IML: Intermediolateral cell column
MLR: Mesencephalic locomotor region
MNs: Motoneurons
ROI: Regions of interest
RF: Reticular formation
RVLM: Rostro-ventrolateral medulla
SPNs: Sympathetic preganglionic neurons
VR: Ventral root

## Data Availability Statements

The raw data supporting the conclusions of this article will be made available by the authors, without undue reservations.

## Ethics Statement

The animal study was approved by the University of Manitoba Bannatyne Campus Animal Care Committee. The study was conducted in accordance with local and institutional requirements.

## Author Contributions

KCC and JWC conceived of the research and wrote the grant applications that supported the research. KCC, JWC and LDR oversaw the research. CVN, NS and LDR performed experiments, LDR, CVN, KCC and JG analyzed data, developed summaries of the findings and developed figures. LDR drafted the initial manuscript and LDR, JWC and KCC wrote and edited the submitted manuscript. All reviewed, edited and critiqued the manuscript.

---

**Supplementary Figure 1.**
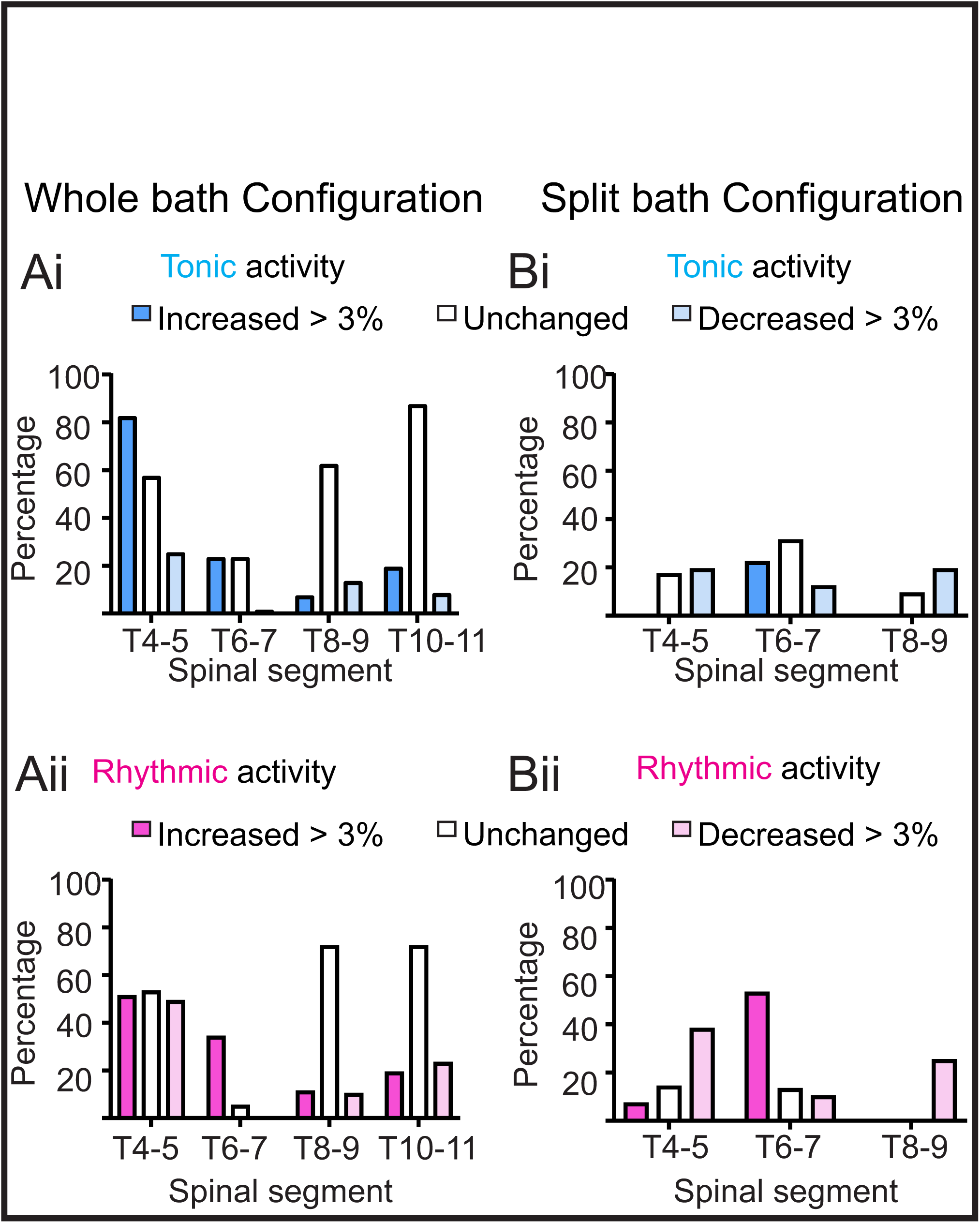
Chi square analysis of segmental distribution of proportions of SPN excitability changes during locomotor activity. Proportions of SPNs demonstrating increases or decreases in Ca-RI, organized by thoracic segment level, during tonic (upper panel, Ai. and Bi.) or rhythmic (lower panel, Aii. and Bii.) activity are not randomly distributed, when organized by thoracic segment level. Chi-square analyses indicated significant differences in all conditions: whole- bath tonic, χ²(6) = 79.47, p < 0.0001; whole-bath rhythmic, χ²(6) = 120.0, p < 0.0001; split-bath tonic, χ²(4) = 37.69, p < 0.0001; split-bath rhythmic, χ²(4) = 84.96, p < 0.0001.

